# Finding Drug Repurposing Candidates for Neurodegenerative Diseases using Zebrafish Behavioral Profiles

**DOI:** 10.1101/2023.09.12.557235

**Authors:** Thaís Del Rosario Hernández, Sayali V Gore, Jill A Kreiling, Robbert Creton

## Abstract

Drug repurposing can accelerate drug development while reducing the cost and risk of toxicity typically associated with de novo drug design. Several disorders lacking pharmacological solutions and exhibiting poor results in clinical trials - such as Alzheimer’s disease (AD) - could benefit from a cost-effective approach to finding new therapeutics. We previously developed a neural network model, Z-LaP Tracker, capable of quantifying behaviors in zebrafish larvae relevant to cognitive function, including activity, reactivity, swimming patterns, and optomotor response in the presence of visual and acoustic stimuli. Using this model, we performed a high-throughput screening of FDA-approved drugs to identify compounds that affect zebrafish larval behavior in a manner consistent with the distinct behavior induced by calcineurin inhibitors. Cyclosporine (CsA) and other calcineurin inhibitors have garnered interest for their potential role in the prevention of AD. We generated behavioral profiles suitable for cluster analysis, through which we identified 64 candidate therapeutics for neurodegenerative disorders.

## Introduction

Alzheimer’s Disease (AD) is the most common cause of dementia among the aging population("2023 Alzheimer’s disease facts and figures," 2023), characterized by senile plaques resulting from accumulation of extracellular amyloid β (Aβ), the formation of intracellular neurofibrillary tangles (NFT) through tau hyperphosphorylation, and overall neuronal degeneration(Anand et al., 2014; Braak & Braak, 1991). Symptoms of AD do not emerge until late in the disease course, obscuring the underlying mechanisms of AD pathology ("2023 Alzheimer’s disease facts and figures," 2023). There is an absence of disease-modifying therapeutics, and current FDA-approved drugs for AD only provide symptomatic relief or marginally prevent the progression of the disease (Mullane & Williams, 2013). Although there is a growing need for both preventative and remedial AD treatments, clinical trials are majorly unsuccessful in remedying AD symptomatology (Mullane & Williams, 2013).

Drug discovery is an extensive and expensive process that often results in compounds that do not make it to market due to safety and/or efficacy concerns, adverse side effects, and incompatibility with the comorbidities present in the target population (Arrowsmith & Harrison, 2012). Drug repurposing is a valuable method to take advantage of the additional targets of currently approved drugs that have already been determined safe for human use. It has been used as a powerful tool to find alternative uses for pharmaceutical compounds that are already on the market, for diseases such as Parkinson’s disease (amantadine), tuberculosis (cycloserine), and attention deficit hyperactivity disorder (atomoxetine)(Al-Subari et al., 2020; Hauser et al., 2017; Roy & Chaguturu, 2017). Cyclosporine A (CsA) is one such drug repurposing candidate in the context of AD. CsA is a calcineurin inhibitor currently used for chronic immunosuppression to prevent allograft rejection (Taglialatela et al., 2015). A human population study found that organ transplant recipients maintained on calcineurin inhibitors, including CsA, had lower incidences of AD compared to the general population (Taglialatela et al., 2015).

We previously developed a deep neural network model Z-LaP Tracker, based on the markerless position estimation software DeepLabCut, to quantify relevant behaviors in zebrafish larvae (Gore et al., 2023). Zebrafish are an established model for the study of a wide variety of pathologies, including neurodegenerative disorders, in part due to their homology with the human genome (Howe et al., 2013; Kalueff et al., 2014). They have comparable behavioral responses to mammalian systems, exhibiting a wide range of behaviors quantifiable by well-established behavioral test batteries (Kalueff et al., 2013; Kalueff et al., 2014; Norton, 2013). In particular, zebrafish larvae are an attractive model for high throughput screens of sizable compound libraries. Small molecule drugs can be easily administered through the water and the larvae can be placed in multi-well plates for phenotypic screening. In the current study, we used a high throughput, whole-organism approach to screen a small-molecule library of FDA-approved drugs and identify CsA-like compounds (Fig 1). We generated a list of potential drug repurposing candidates for the prevention and treatment of neurological disorders, including AD.

## Results

### Behavioral Screening of FDA-Approved Drugs

Zebrafish larvae were exposed to the 876 compounds included in the Cayman Chemical FDA-approved Drug Library for a total of 6 hours. Each compound was evaluated using 48 zebrafish larvae, and each experiment included DMSO and egg water controls. In total, we examined 50,496 larvae: 42,048 larvae treated with small-molecule compounds, and 8,448 DMSO-treated larvae. Zebrafish larvae were exposed to all compounds and controls at a concentration of 10μM for 6 hours. At this concentration 23 compounds resulted in a ≥50% mortality rate, but none of the compounds had a 100% mortality rate. We did not include larvae if they moved less than 1% of the time throughout the experiment. Additionally, low likelihood (< 0.50) data points were filtered out and not included in behavioral profile generation. We generated behavioral profiles by comparing differences in behavior between DMSO-vehicle controls and each compound.

Our behavioral assay features a 3-hour Microsoft PowerPoint presentation with an initial 60 minutes of no stimuli, 80 minutes of moving lines - alternating direction every 10 minutes and color every 20 minutes -, followed by a 10 minute period with no stimuli, and 30 minutes of sound stimuli alternating frequency every 10 minutes (Fig 2A). This assay is analyzed in 10-minute intervals called periods, and behaviors are averaged per period.

**Fig. 2.**
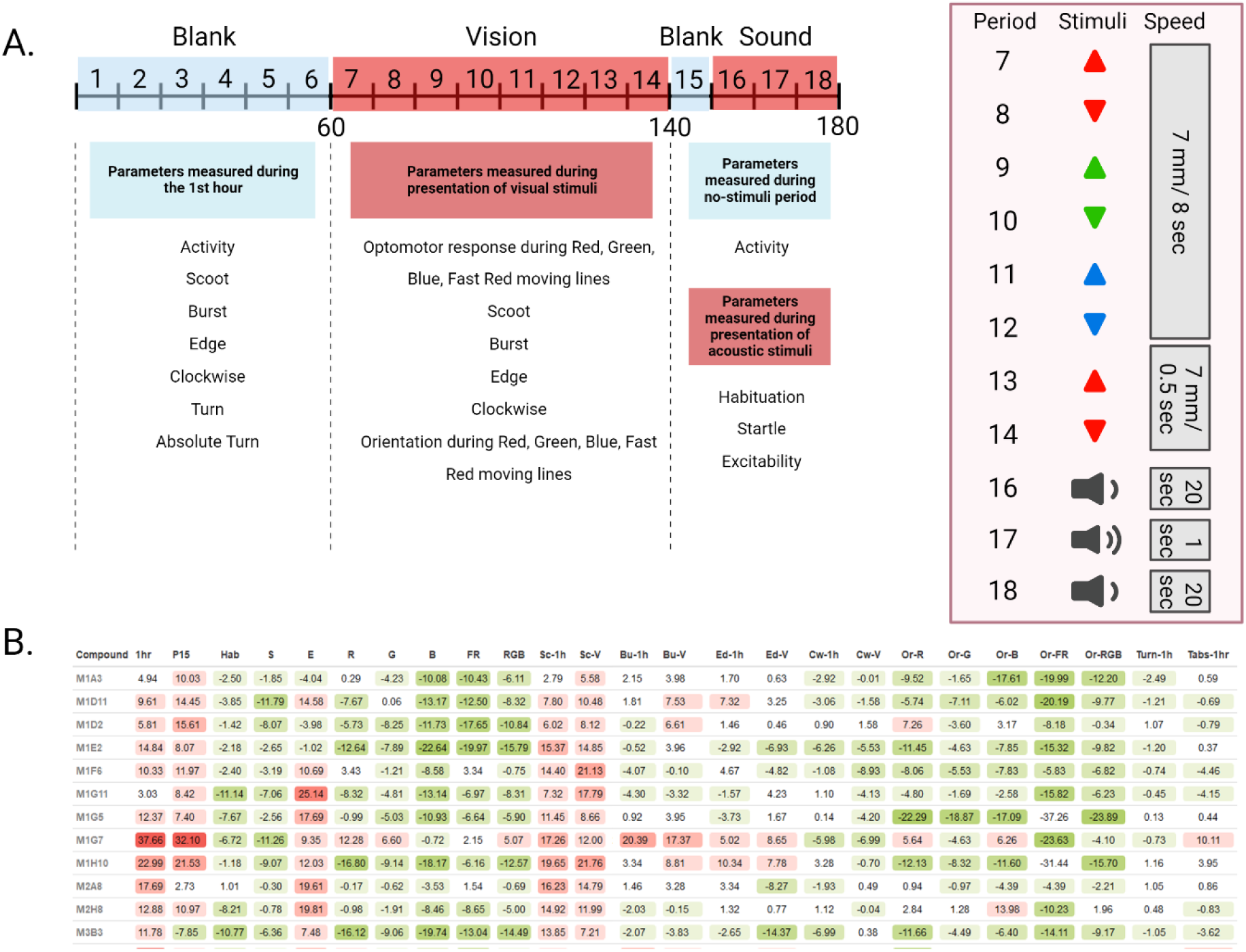
Overview of 25 parameters measured during 3-hour behavioral assay. **(A)** 3-hour timeline of the presentation shown to treated zebrafish larvae. 25 parameters of behavior are measured during 18 periods of visual and acoustic stimuli. **(B)** Behavioral profiles of treatments using 12 compounds from the Cayman Chemical FDA-approved Drug Library. Each treatment is compared to DMSO-vehicle controls and the resulting differences are color-coded based on an increase (red) or decrease (green) in a particular behavioral parameter. The values represent differences as compared to DMSO controls in percentage points. Treatments are identified by the Cayman Chemical library’s plate number (M1-M11) and well number (A1-H11).

The resulting behavioral profiles consisted of 25 behaviors representing overall activity, reactivity, swimming patterns, and optomotor response (Fig 2B). Namely, we measured (1) Activity during the 1st hour, (2) Activity during Period 15, (3) Habituation, (4) Startle response, (5) Excitability, (6) Optomotor response to moving red lines, (7) Optomotor response to moving green lines, (8) Optomotor response to moving blue lines, (9) Optomotor response to faster moving red lines, (10) Combined optomotor response to red, green, and blue moving lines, (11) Scoot movement during the 1st hour, (12) Scoot movement during the presentation of moving lines of any color or speed, (13) Burst movement during the first hour, (14) Burst movement during the presentation of moving lines of any color or speed, (15) Percent edge location during the first hour, (16) Percent edge location during the presentation of moving lines of any color or speed, (17) Percent clockwise orientation during the 1st hour, (18) Percent clockwise orientation during the presentation of moving lines of any color or speed, (19) Upward orientation during moving red lines, (20) Upward orientation during moving green lines, (21) Upward orientation during moving blue lines, (22) Upward orientation during faster moving red lines, (23) Combined upward orientation during red, green, and blue moving lines, (24) Turn angle during the 1st hour, and (25) Absolute turn angle during the 1st hour (Supplementary Table 1).

## Clustering Results

### K-means Cluster Analysis

We used the K-means clustering algorithm to identify compounds in the Cayman Chemical FDA-approved Drug Library that would cluster together with calcineurin-inhibitor CsA. We identified the optimal number of clusters (k = 4) using the elbow method. We found that CsA clusters with 58 other compounds (Fig 3). The overall behavioral profile of CsA-like compounds features increased activity during the first hour and period 15, decreased habituation and startle response, increased excitability, decreased optomotor response during all visual stimuli, increased Scoot movement, and a reduced orientation response during all visual stimuli (Fig 4). The Pearson correlation value for this cluster was 0.62.

**Fig. 3.**
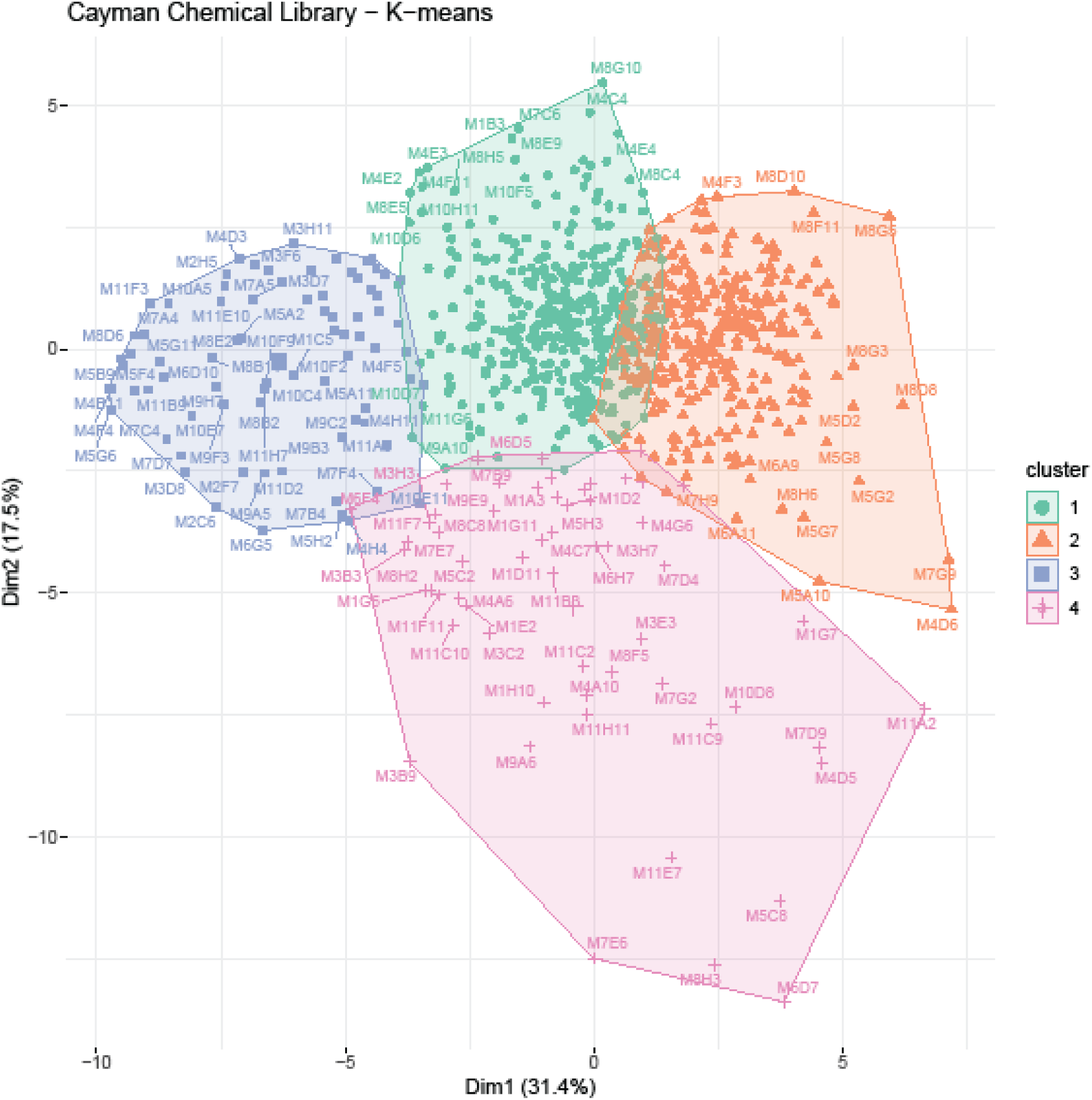
K-means cluster analysis. 876 compound treatments and DMSO controls were assigned to k=4 clusters. Cluster 4 contains cyclosporine A and 58 compound treatments with similar behavioral profiles.

**Fig. 4.**
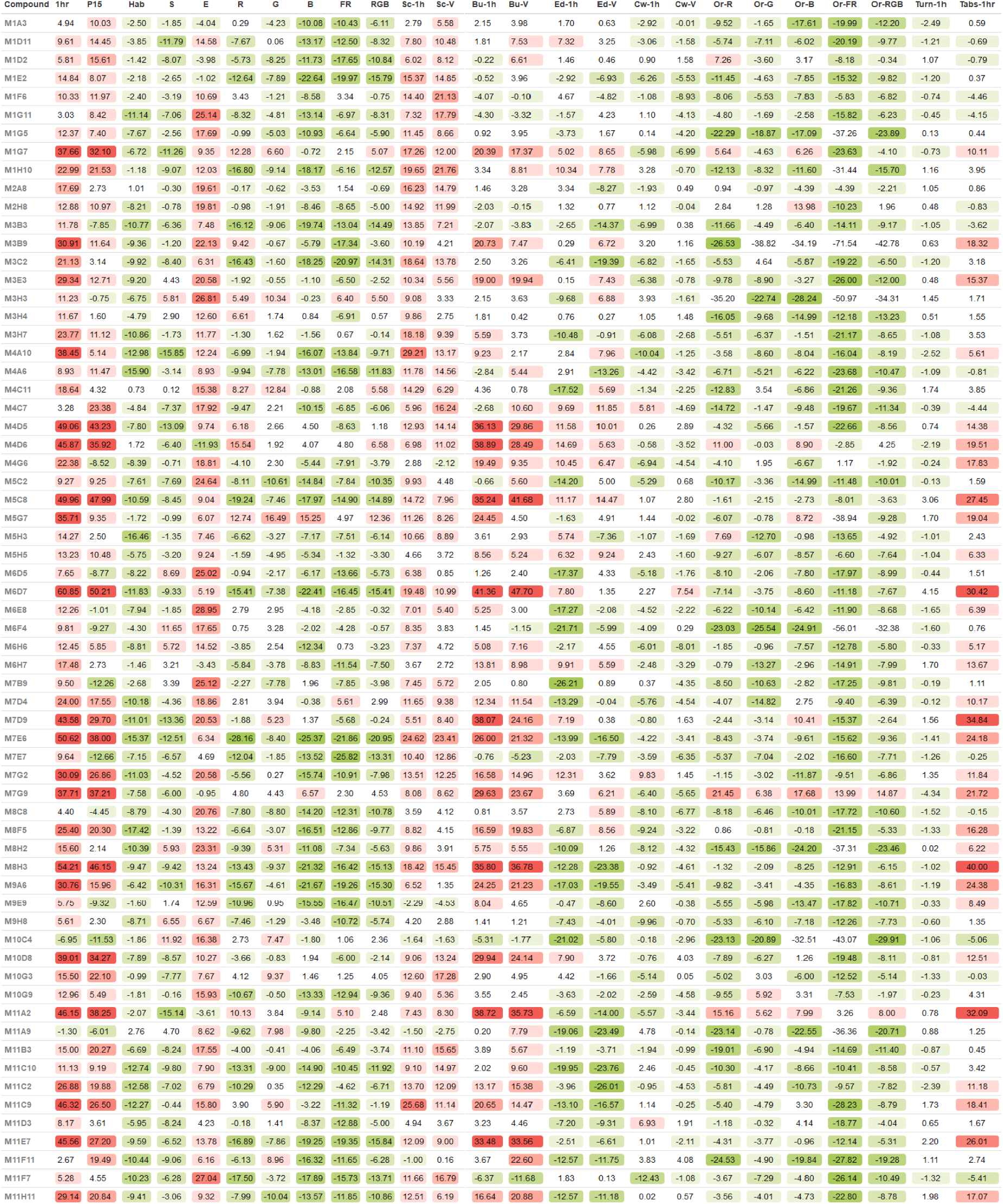
Behavioral profiles of CsA-like compounds. Identified 64 compounds that induce behavioral profiles similar to CsA when administered to 5 dpf zebrafish larvae. Each behavioral profile is composed of 25 parameters measuring activity, reactivity, swimming patterns, and optomotor response.

### Hierarchical Cluster Analysis

We used agglomerative hierarchical clustering to group the behavioral profiles generated by exposure to the Cayman Chemical FDA-approved Drug Library. We found that CsA and other compounds with similar behavioral profiles formed a distinct cluster featuring the same behavioral patterns found in our previous screening of FDA-approved drugs (Fig 5A) (Tucker Edmister, Del Rosario Hernández, et al., 2022). We identified 53 CsA-like compounds, of which 47 were also found with the K-means clustering method (Fig 5B). The correlation value for this cluster was 0.60. Multiple CsA-like subclusters with increasing correlation values can be identified in our hierarchical analysis (Supplementary Figure 1).

**Fig. 5.**
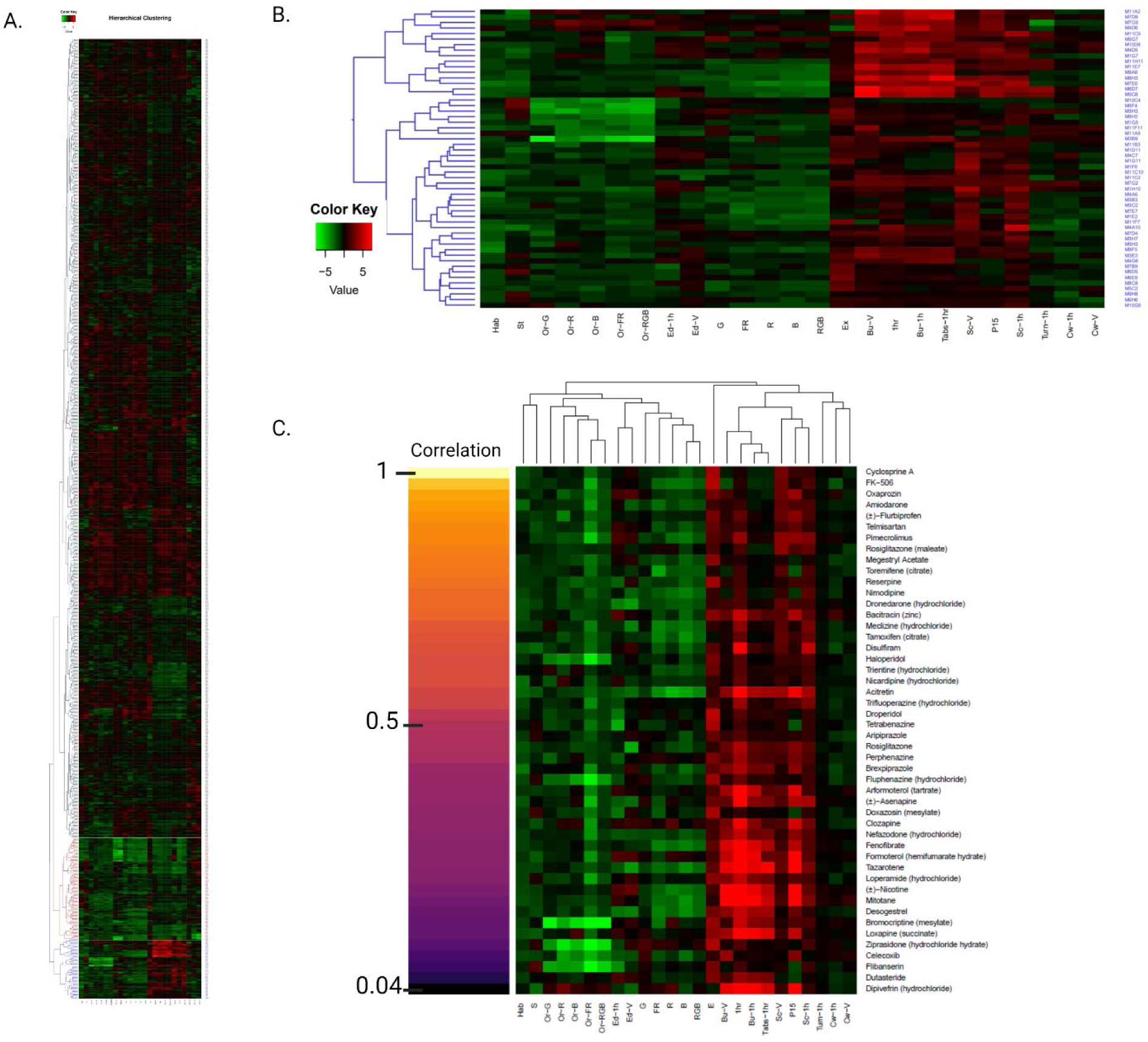
Hierarchical cluster analysis. **(A)** Overview of the behavioral profiles elicited by 876 compound treatments. A high-resolution version of this image is included in the supplementary information (Supplementary Figure 2). **(B)** Cluster of 53 compounds that induce CsA-like behavioral profiles. Red indicates an increase in a behavioral value relative to DMSO controls, while green indicates a decrease. **(C)** Pearson pairwise correlations of CsA and the 47 CsA-like compounds identified by both K-means and hierarchical clustering.

Using K-means and hierarchical cluster analysis, we found a total of 64 compounds displaying CsA-like behavioral paradigms (Table 1). These 64 compounds affect manifold biological functions and are used to treat a wide range of diseases. We further analyzed the composition of our clusters of interest by calculating the Pearson correlation between the behavioral profiles of the compounds in the Cayman Chemical FDA-approved Drug Library and CsA. We visualized the 47 compounds found by both clustering methods showing their degree of similarity to CsA (Fig 5C).

**Table 1.**
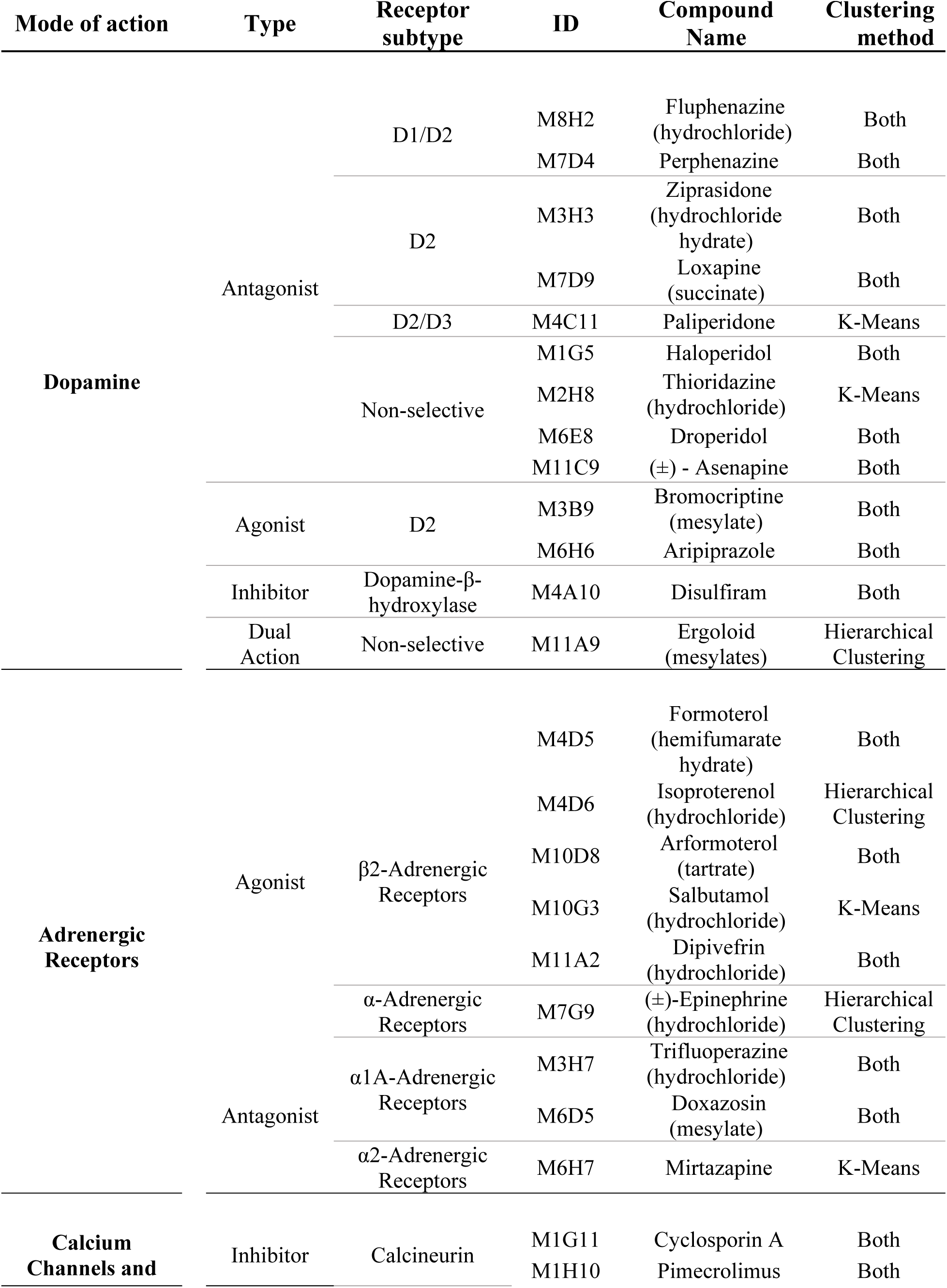
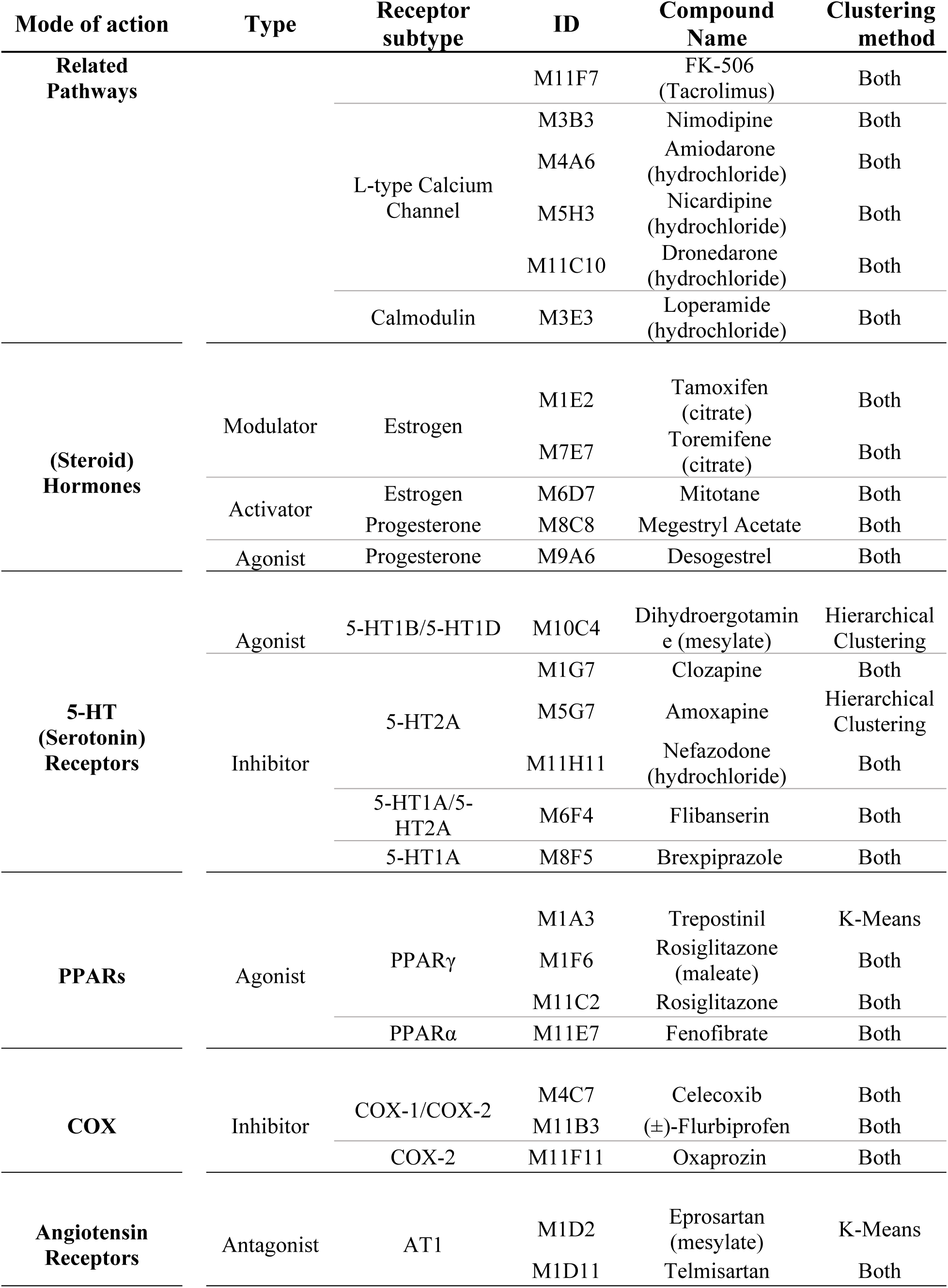
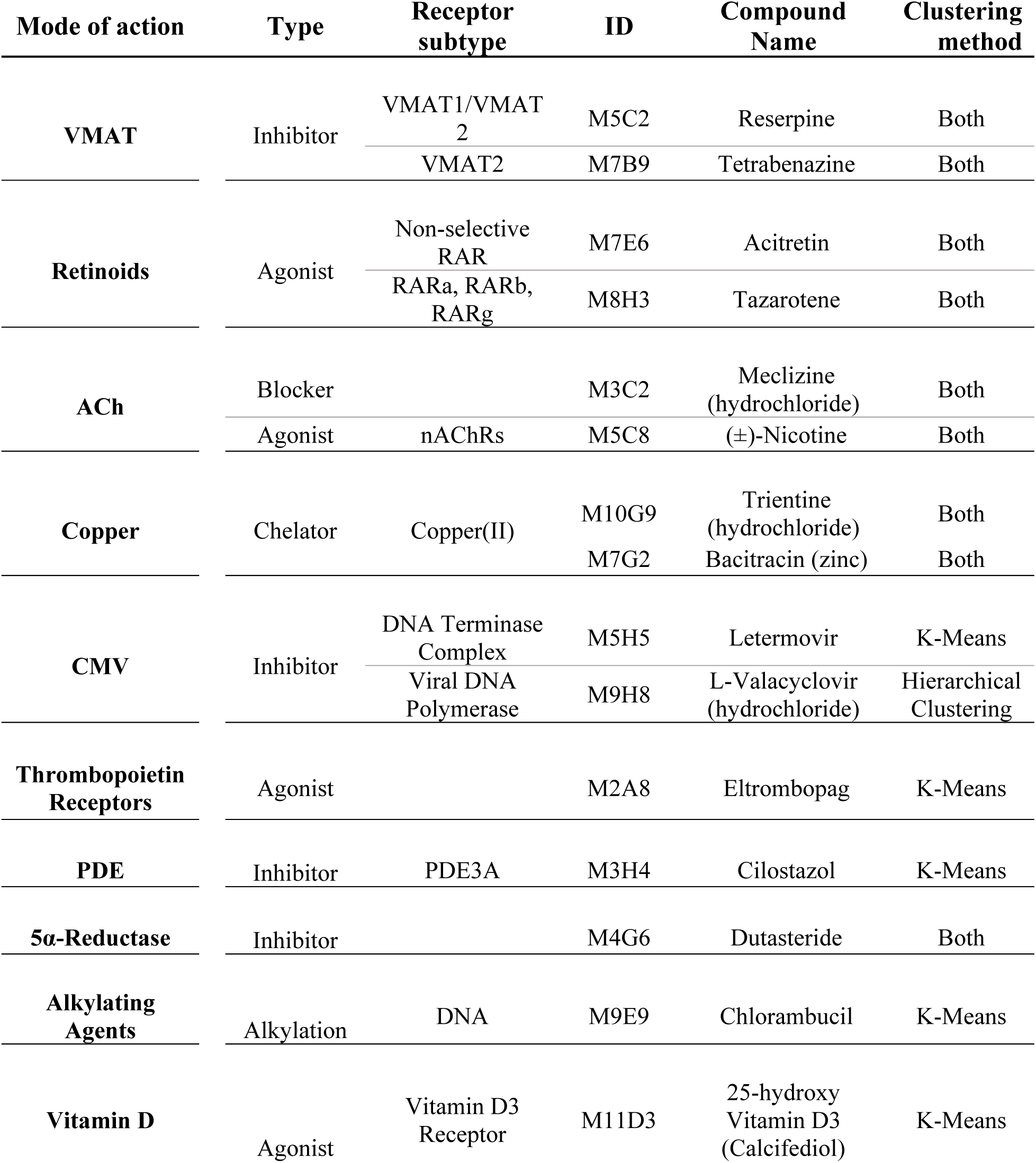
List of compounds inducing CsA-like behavioral profiles, identified by K-means and hierarchical clustering analyses.

### Novel Compounds Displaying CsA-like Behavior

Our K-means clustering and Hierarchical clustering analyses revealed a total of 64 compounds with behavioral paradigms similar to CsA. Of these, 47 were found in both cluster analysis methodologies. We performed statistical analyses on the effects of these compounds in larval behavior. When compared against DMSO-vehicle controls, 89% of the CsA-like compounds induced statistically significant changes in behavior. These changes were observed throughout most of the 25 behavioral measures, excluding clockwise movements and turn angle. Overall, 40% of the compounds screened in this study elicited at least one significantly different behavior compared to DMSO controls. The statistical analyses performed on the behavioral profiles of all the compounds found in the Cayman Chemical FDA-approved Drug Library can be found in Supplementary Table 2.

### Identification of Predominant Targets and Pathways in CsA-like Clusters

To further investigate potential similarities between the identified CsA-like compounds, we queried the Disease-Gene Interaction Database (DGIdb), Therapeutic Targets Database (TTD), Guide to Pharmacology (GtoPdb), Kyoto Encyclopedia of Genes (KEGG), Protein ANalysis THrough Evolutionary Relationships (PANTHER), WikiPathways, and Reactome databases and matched each compound in the Cayman Chemicals Library of FDA-approved Drugs to primary and secondary target genes, as well as their respective molecular pathways and associated mechanisms of action. Our multi-database exploration approach enabled us to fill in information gaps between sources and resulted in an unbiased collection of information, which would otherwise not be possible without an extensive literature review.

We compared the most common molecular targets perturbed by the cluster of CsA-like compounds to the overall target composition of all the compounds in the Cayman Chemical FDA-approved Drug Library. We found that the CsA-like cluster compounds acted on an entirely different composition of targets than the predominant targets found in the compilation of all compounds in the small-molecule library. Specifically, the most common targets of the entire library were genes encoding P450 enzymes (Fig 6A). In contrast, the CsA-like drugs predominantly target genes related to neuromodulation (dopamine receptors, serotonin receptors, adrenergic receptors) or associated with neurodegenerative disorders (ATXN2, KCNH2, mTOR) (Fig 6B).

**Fig. 6.**
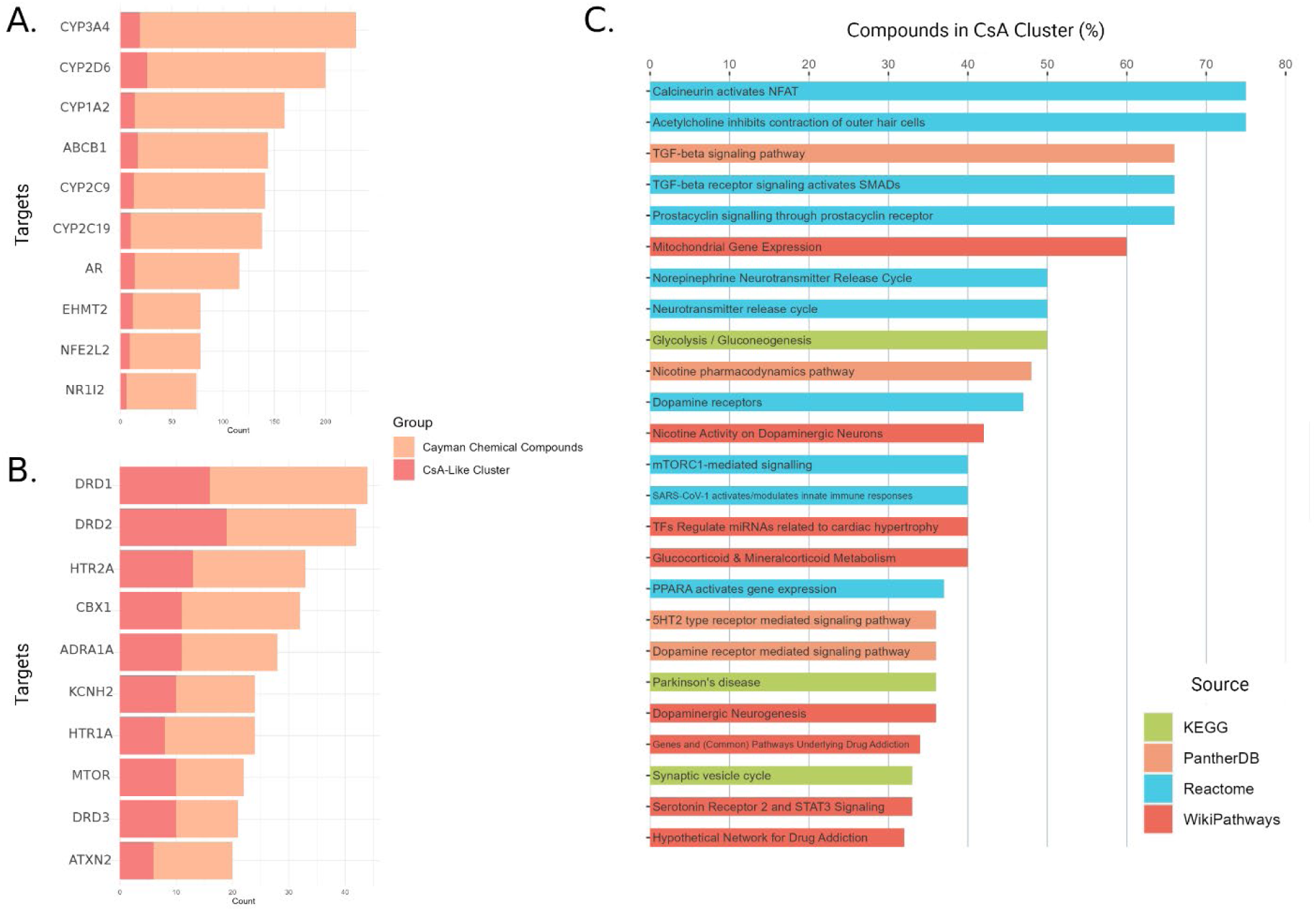
Predominant targets and pathways perturbed by CsA-like compounds. **(A)** Top 10 targets affected by the 876 compounds in the Cayman Chemical FDA-approved Drug Library. **(B)** Top 10 targets affected by CsA and the 64 CsA-like compounds found in our clustering analyses. **(C)** Top 25 Wikipathways, Reactome, KEGG, and PANTHER pathways containing targets affected by CsA-like compounds.

We queried all the targets affected by the 64 CsA-like compounds and matched them to biological pathways from Wikipathways, Reactome, KEGG, and PANTHER (Fig 6C). We selected the top 25 pathways by percentage of CsA-like compounds compared to other compounds not found in our clustering analyses. The pathways’ categories correspond with the predominant targets from our previous query, continuing to demonstrate an emphasis on neurological function and modulation.

Additionally, we utilized Ingenuity Pathway Analysis (IPA) to map the predicted relationships between CsA-like compounds and AD-related targets. We generated a custom pathway and overlaid connections with the Molecule Activity Prediction (MAP) tool available in the IPA software (Fig 7). The resulting network predicts a heavily inhibited AD node through diverse downstream effects triggered by the activation of CsA-like molecules.

**Fig. 7.**
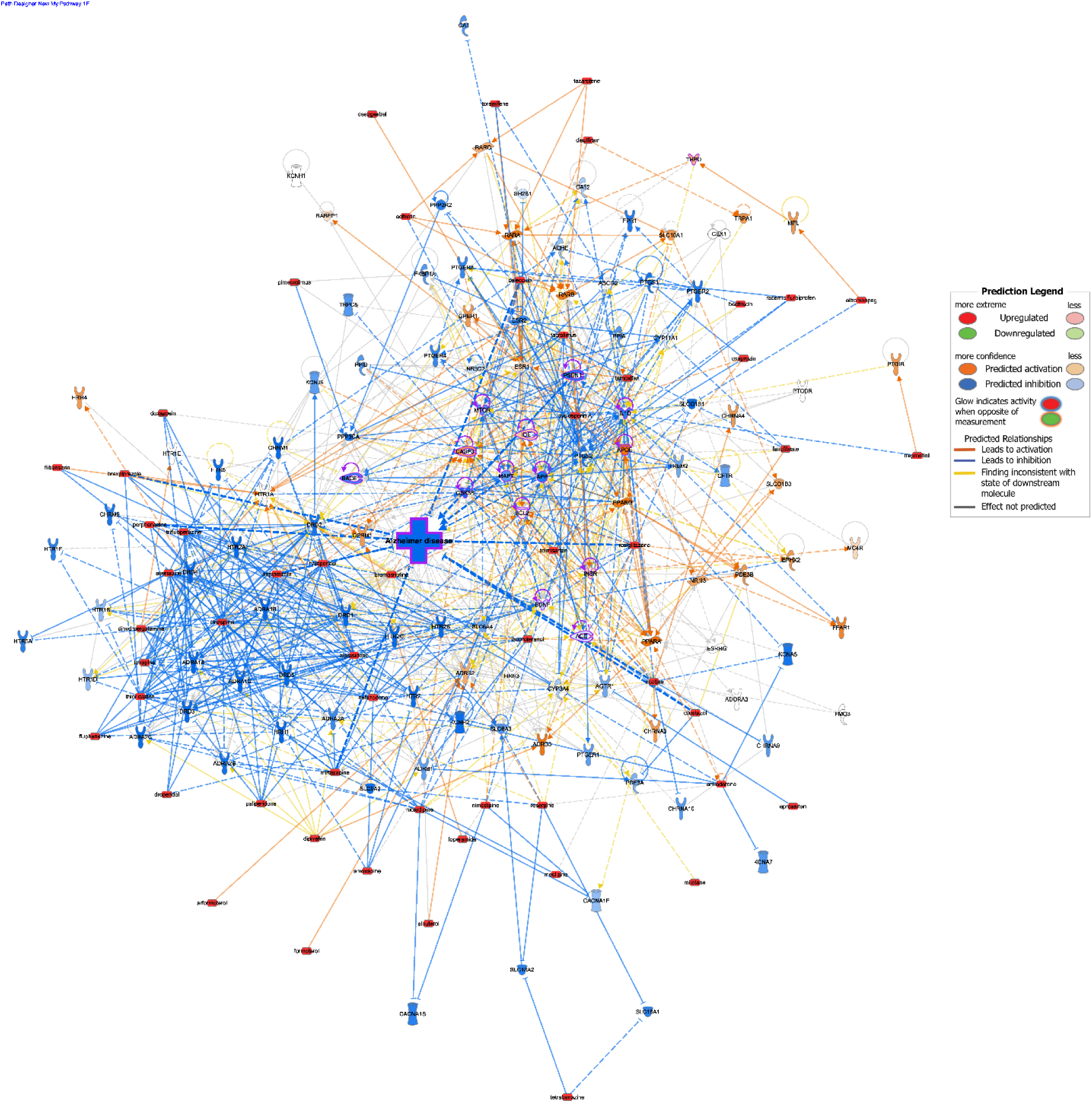
IPA analysis showing the predicted effect of CsA-like compounds on Alzheimer’s disease. Orange lines and nodes indicate predicted activation, blue lines and nodes indicate predicted inhibition, and yellow lines indicate inconsistent findings. Red nodes indicate activation through custom input using the Molecule Activity Predictor (MAP). Purple outlines indicate AD-related nodes.

## Discussion

In this study, we evaluated 876 FDA-approved compounds in 5dpf zebrafish larvae. We generated behavioral paradigms for each compound and clustered them to find CsA-like compounds. Using K-means and hierarchical clustering, we found a total of 64 compounds that evoke CsA-like behavior in 5 dpf zebrafish larvae. We chose to use two clustering analyses to classify our behavioral profiles. Due to the large number of measurements obtained in the current study, we sought to reduce the number of volatile factors during our analysis - factors such as overrepresentation of behaviors and outliers within the data could all have an unexpected effect on clustering results. We employed and compared the results of both clustering methods to maximize the generation of recognizably stable patterns from our behavioral data. Since each method employs different criteria to cluster the behavioral profiles of FDA-approved drugs, they serve as mutual confirmation of their results - evidenced by the high amount of overlap between the identified clusters. They also provide us with more conservative measures to evaluate the produced clusters, such as a list of compounds whose patterns persisted through clustering methods.

In a previous study, we screened 190 compounds included in the Tocriscreen FDA-Approved Drugs Library using the same methods described in the current study and found 32 compounds displaying CsA-like behavior (Gore et al., 2023). These compounds act on a variety of molecular targets, pathways, and diseases, yet induce analogous patterns in zebrafish larval behavior during our 25-behavior screening. During our current screening of 876 compounds, we found a total of 64 compounds displaying CsA-like behavior, of which 11 were previously found in our previous screening, and 2 were not present during our current screening. There are 19 compounds that were included in both screenings, but only found to cluster with CsA during our previous screening. There is a majority but not a complete overlap between the compounds found in the Tocriscreen and the Cayman Chemical libraries, given that a total of 172 compounds are shared between the two libraries. This gives rise to some questions about the reproducibility of the “missing” compounds - whether they truly display a CsA-like behavioral profile. A possible explanation may be that some of these compounds display behavioral profiles akin to other compounds within their target group, which is made more evident in a larger screening and results in tighter biological classification-based clusters, and the rest induce behaviors highly deviated from the range typically found in CsA-like compounds (Supplementary Figure 3).

We classified the 64 CsA-like compounds found during our clustering analyses and noticed that they targeted 5 main categories: dopamine receptors, adrenergic receptors, calcium channels and related pathways, steroid hormones, and 5-HT receptors. To further investigate the clinical significance of our results, we manually curated literature pertaining to these 64 compounds in association with AD or Alzheimer’s-like pathology (Table 2).

**Table 2.**
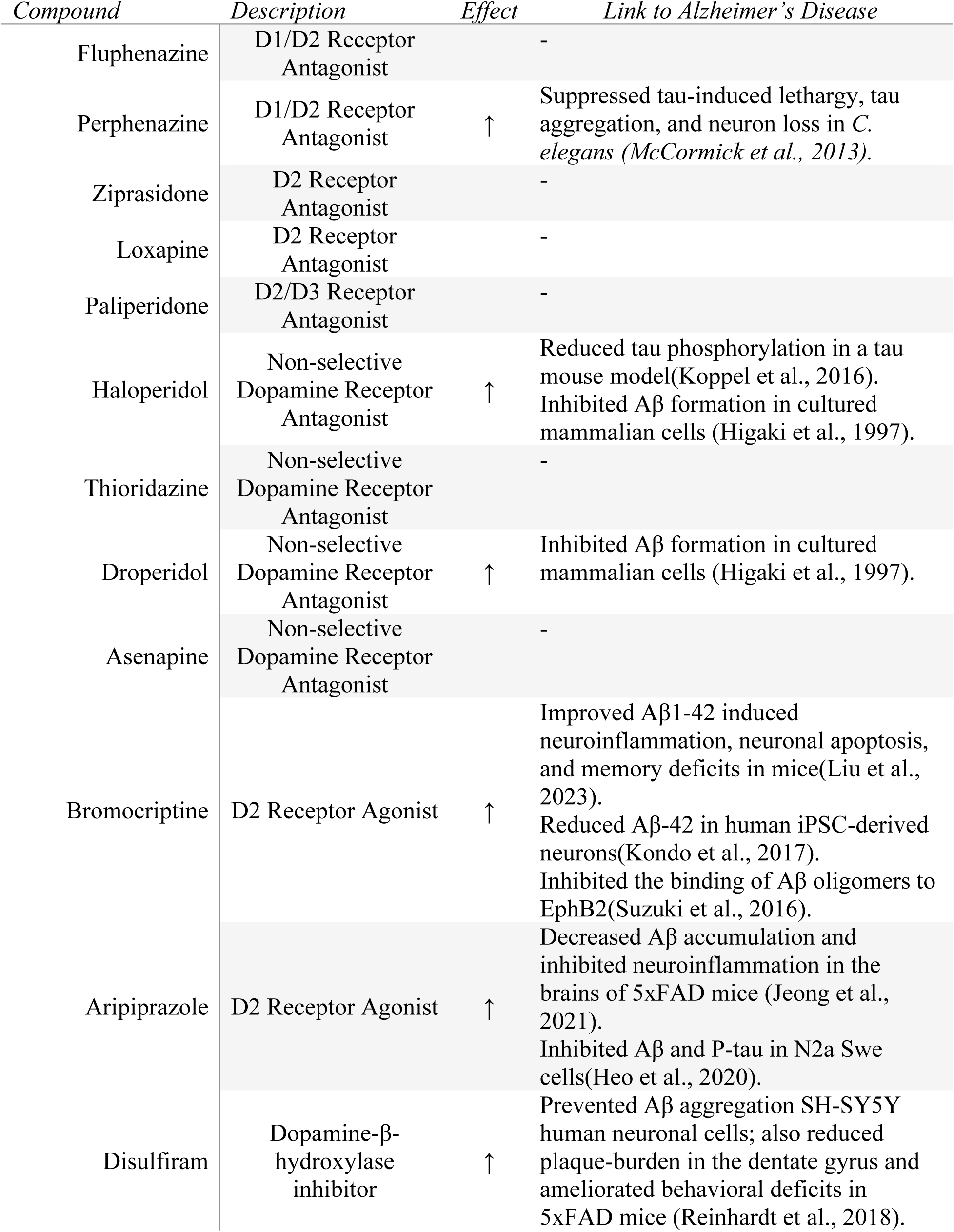
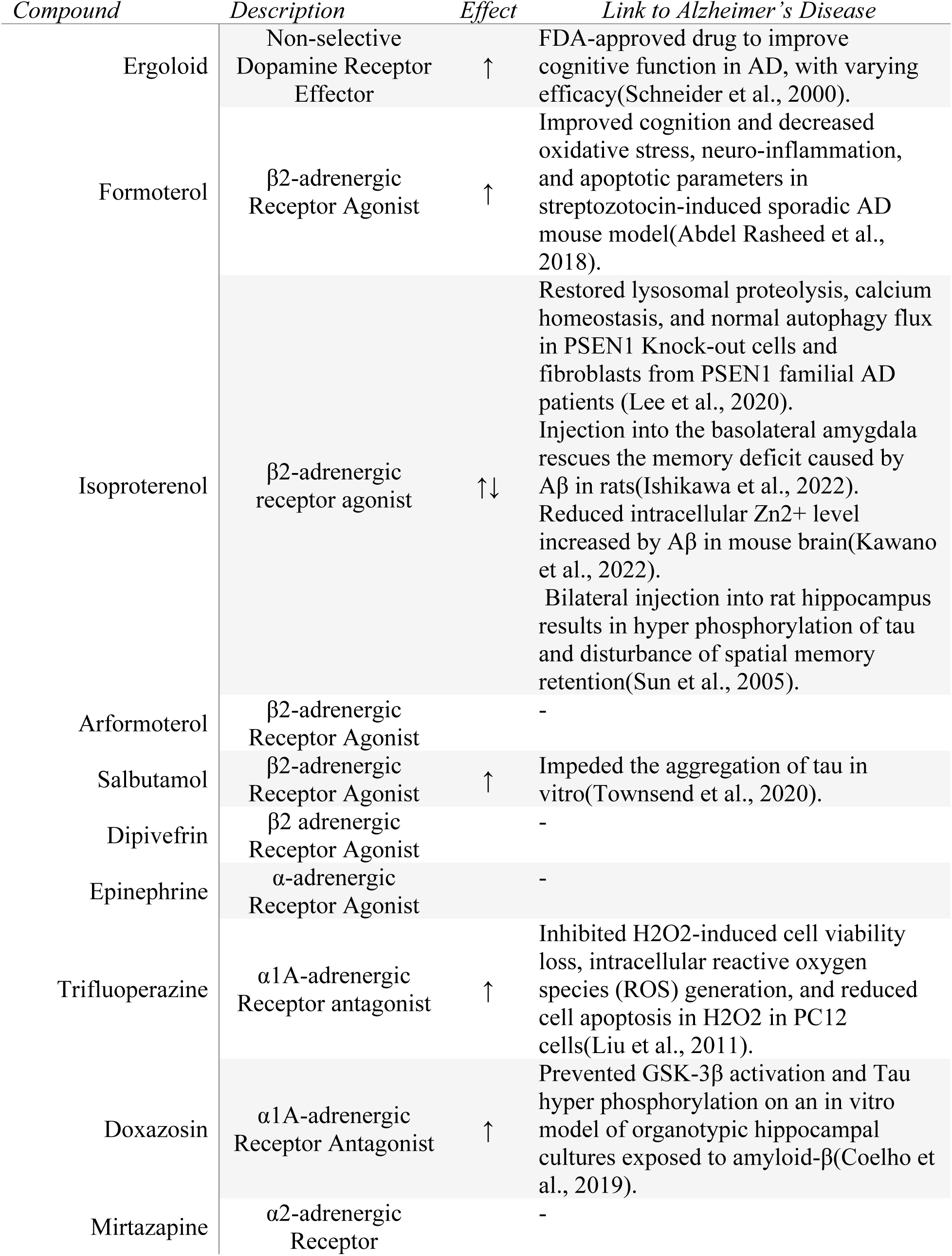
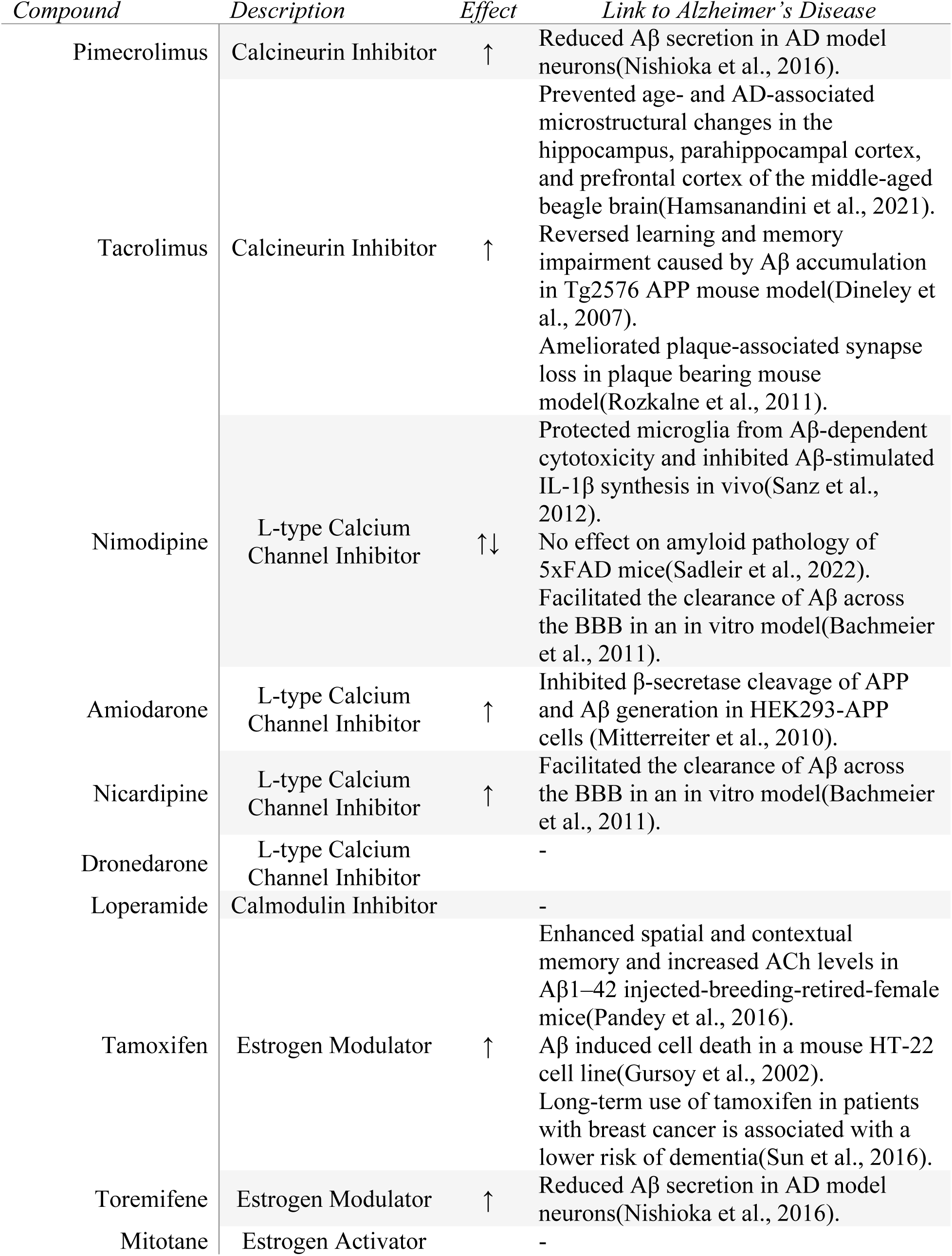
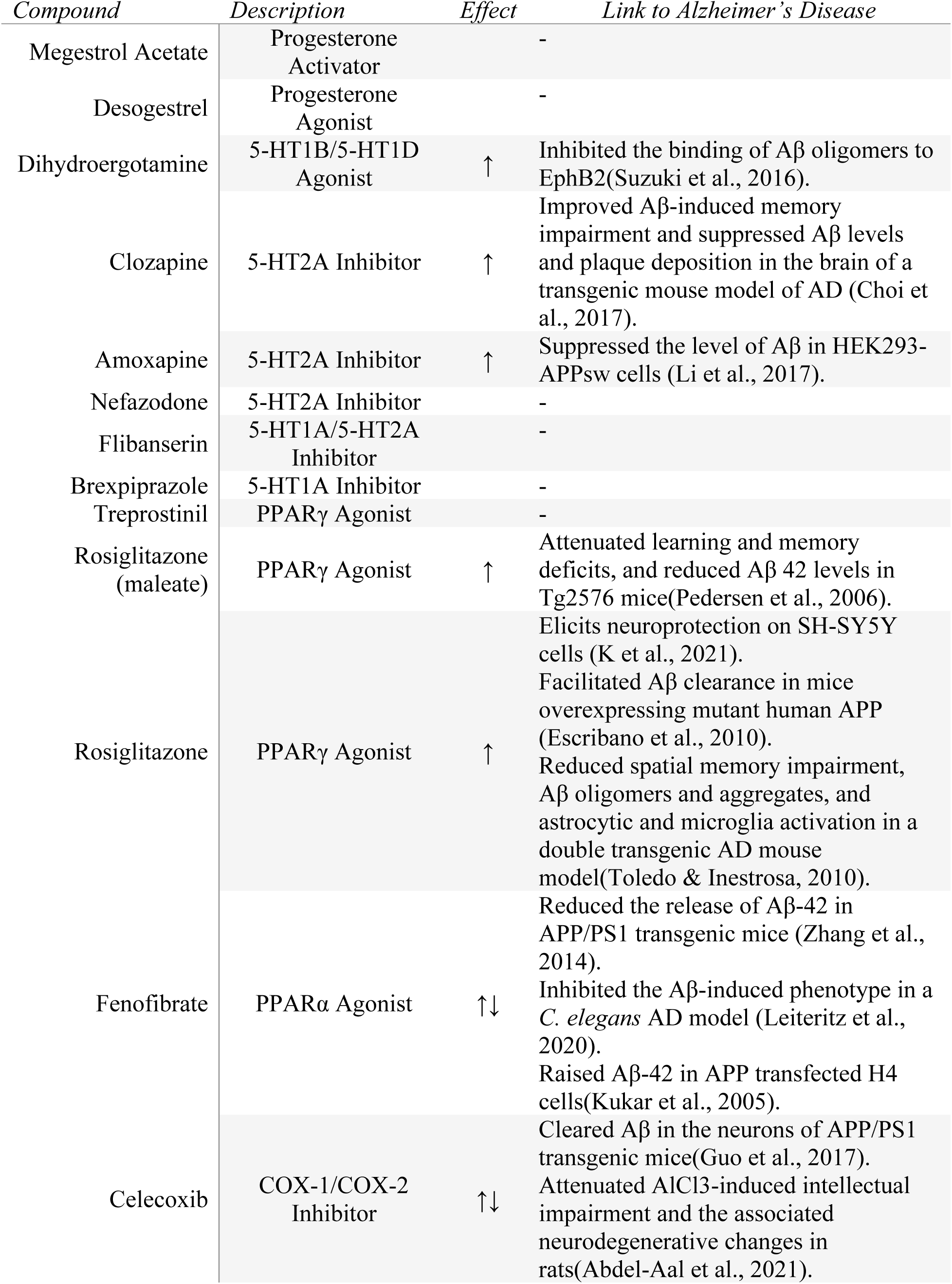
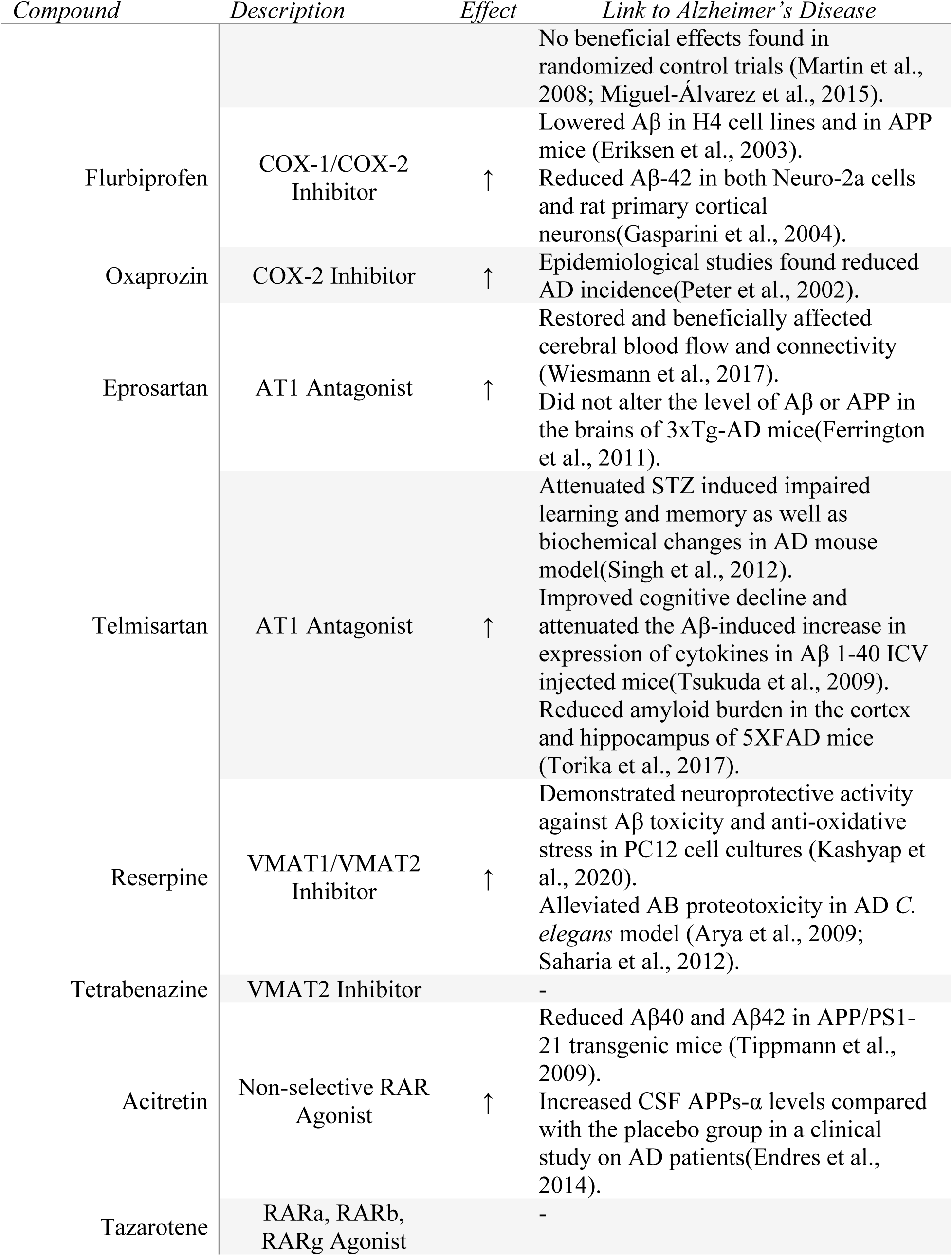
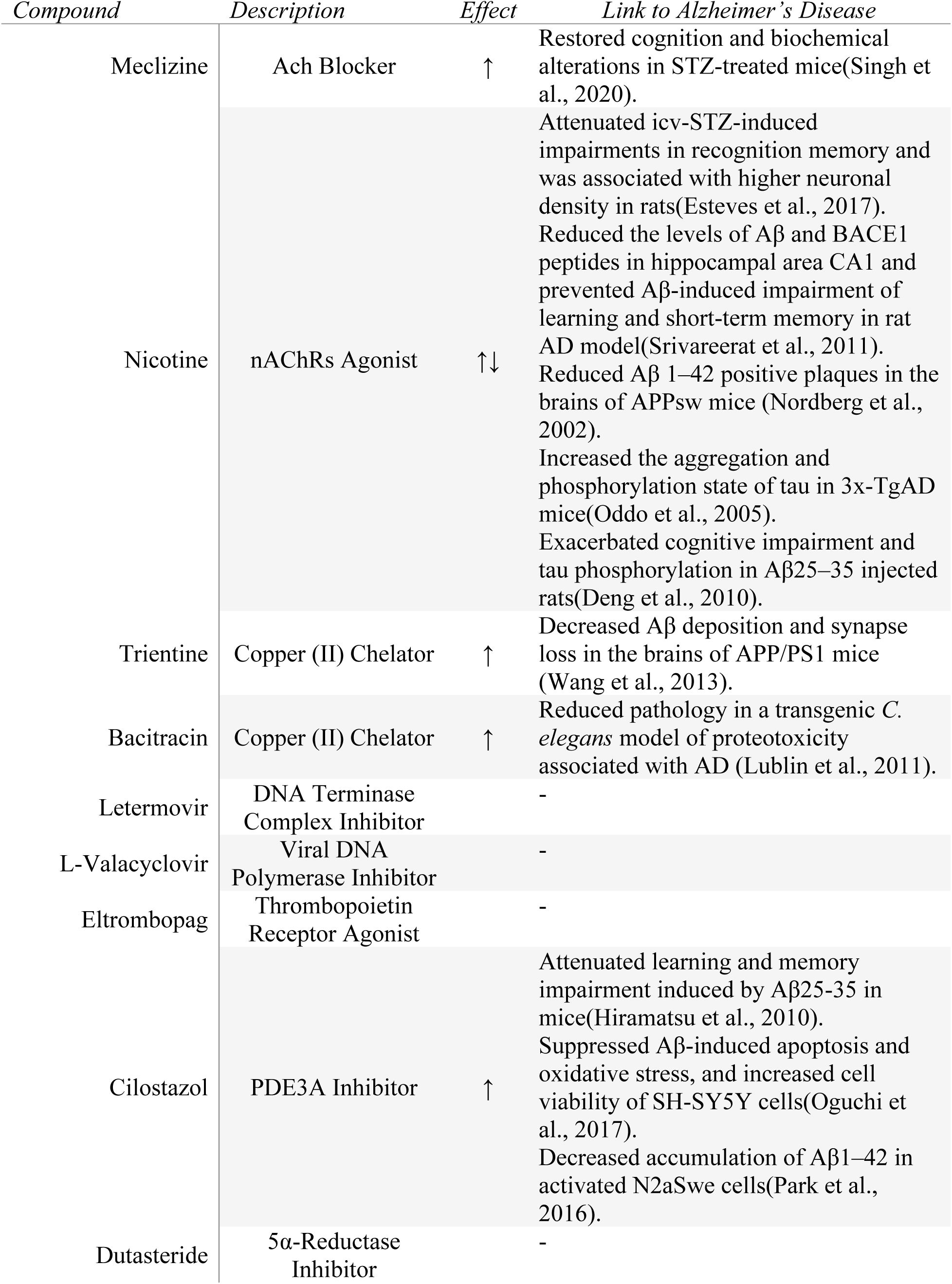
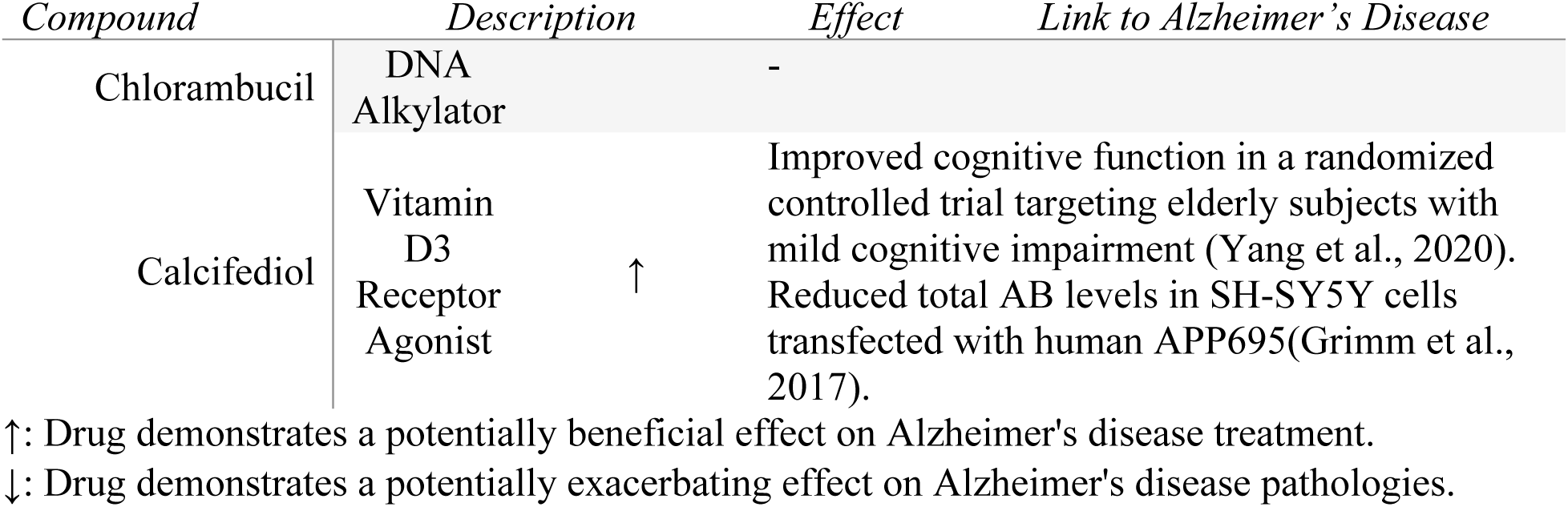
Literature search for CsA-like compounds’ association with Alzheimer’s disease.

Dopaminergic system dysfunction has been associated with AD through multiple post-mortem and in vivo studies measuring dopamine levels, cortical plasticity, and dopaminergic neuron degeneration (D’Amelio et al., 2018; Kemppainen et al., 2003; Nobili et al., 2017; Storga et al., 1996). These findings propose a strong link between dopaminergic deficit and AD in both early and late stages of the disease. A study using dopamine agonist rotigotine found an increase in cholinergic activity and normalized levels of LTP-like cortical plasticity (Koch et al., 2014). We found a total of 9 dopamine receptor antagonists and 2 dopamine receptor agonists of interest during our screening. The majority of the dopamine receptor antagonists in our list have not been studied in the context of AD; however, some dopamine receptor antagonists - including Haloperidol - have been chosen for clinical trials (D’Amelio et al., 2018). Furthermore, the 4 remaining dopamine-associated compounds found in our CsA-like cluster have all been linked to AD pathology, mostly through the inhibition of Aβ aggregation and improvement of cognitive function.

Dopamine-related antipsychotics, both typical and atypical, are the primary pharmacological option used to treat the neuropsychiatric and behavioral symptoms often present in AD (Calsolaro et al., 2019). These can include agitation, depression, psychosis, and overall behavioral disturbance. Several of the compounds listed in the present study are used to treat these symptoms in AD patients, namely, fluphenazine (Gottlieb et al., 1988), perphenazine (Pollock et al., 2002), ziprasidone (Kasckow et al., 2004), loxapine (Petrie et al., 1982; Raskind, 1998), and haloperidol (Devanand et al., 1998). It is important to note that many of these compounds have multiple targets and can simultaneously affect a variety of pathways. For instance, there is evidence of interaction between the dopamine and the serotonin pathways, implying the possibility of cross-pathway alteration in AD pathology (Ceyzériat et al., 2021). This is specifically utilized in the prescription of atypical antipsychotics, distinctively known as serotonin-dopamine antagonists. Due to their relevant multi-target capabilities and current use in patients with AD, we believe that this class of drugs would benefit from further studies exploring their possible benefits in the treatment of multiple stages of AD progression.

β-adrenergic receptors (βARs) are G protein-coupled receptors (GPCRs) responsible for regulating synaptic plasticity and memory formation(Kemp & Manahan-Vaughan, 2008). Specifically, β2-adrenergic receptor (β2AR) agonists are emerging therapeutic targets for neurological diseases (Li, 2023). During our screening, we found 5 β2AR agonists of interest with varying levels of supporting literature in the context of AD. Interestingly, there has been evidence of a reduction in AD prevalence in patients prescribed with β-blockers to treat hypertension (Khachaturian et al., 2006; Rosenberg et al., 2008). This protective trend also applies to other antihypertensive drugs (Ding et al., 2020; Wei et al., 2015), which include α1-adrenergic receptor antagonists such as Doxazosin. However, clinical studies often show contradictory results regarding the effects of β-blockers on cognitive impairment (Lebouvier et al., 2020). Studies performed in AD animal models also highlight the conflicting nature of β2AR signaling in the context of AD (Branca et al., 2014; Ni et al., 2006). Due to the opposing effects of β1- and β2-blockers in memory and cognitive function (Ramos et al., 2005; Ramos et al., 2008), it is difficult to predict their effectiveness in treating AD pathogenesis. Comparably, α-adrenergic receptors (αARs) have been extensively linked to cognition as well as glucose metabolism - both part of AD symptomatology (Gannon et al., 2015). αAR antagonists, in particular, have been investigated both in vitro and in vivo as promising therapeutic targets of AD(Karczewski et al., 2018; Katsouri et al., 2013; Yu et al., 2022), and could prove interesting targets for future experiments.

The potential role of calcium homeostasis dysregulation in AD has been broadly explored through the impairment of mitochondrial function in the presence of excessive calcium levels (Calvo-Rodriguez & Bacskai, 2021), the induction of mitochondrial calcium overload by Aβ oligomers (Calvo-Rodríguez et al., 2016; Sanz-Blasco et al., 2008), and the direct effect of insoluble tau on calcium dyshomeostasis (Esteras et al., 2021; Mahoney et al., 2020). L-type calcium channels are substantially expressed on neurons (Zamponi, 2016), thus making L-type calcium channel blockers such as nimodipine, amiodarone, nicardipine, and dronedarone attractive targets for calcium regulation in the context of AD. Lebouvier et al. outline three potential mechanisms of action of L-type calcium channel blockers: (i) suppression of Aβ-induced calcium release, (ii) inhibition of amyloidogenesis, and (iii) increasing Aβ transcytosis (Lebouvier et al., 2020). Given our initial interest in calcineurin inhibitor CsA, we also emphasize both Pimecrolimus and Tacrolimus as compounds of interest due to both their shared mechanism of action and comprehensive supporting research on the subject of neurodegenerative diseases.

The primary steroid hormones estrogen, progesterone, and testosterone can be found in various regions of the central nervous system (Garcia-Ovejero et al., 2005). Steroidogenesis has been proven to be a natural mechanism to combat neurodegenerative conditions, and in vivo studies have shown an increase in AD-like neuropathology following gonadectomy (Carroll & Rosario, 2012). While hormone therapy seems promising as a preventive therapy in neurodegeneration, epidemiological studies and clinical trials reveal controversial results (Henderson, 2014). Treatment timing and dosage are important factors to consider in future research involving these agents.

The serotonergic system has been associated with cognitive function and performance in neurological diseases including schizophrenia, epilepsy, and AD (Švob Štrac et al., 2016). Neurotransmitter serotonin (5-HT) is involved in the regulation of various physiological processes including cognition and emotional behavior (Ciranna, 2006). Cerebrospinal fluid 5-HT levels in AD patients were significantly decreased compared to healthy controls (Tohgi et al., 1992), and post mortem studies show a decrease in brain 5-HT levels (Palmer et al., 1987). Given the overwhelming evidence of serotonergic influence in cognitive function, there has been interest in various 5-HT receptor (5-HTR) agonists and antagonists for the treatment of AD, including 5-HT2A (Garcia-Romeu et al., 2022; Lu et al., 2021) and 5-HT6 (Geldenhuys & Van der Schyf, 2009; Kucwaj-Brysz et al., 2021). We have identified 6 5-HTR antagonists with various 5-HT receptor targets: dihydroergotamine, clozapine, amoxapine, nefazodone, flibanserin, and brexpiprazole. Considering that multiple 5-HTRs have a demonstrated beneficial effect on cognitive processes, a multi-receptor approach through one or more compounds might be valuable in future studies.

In addition to the aforementioned 5 categories, we also found compounds belonging to 13 other classes, namely, peroxisome proliferator-activated receptor (PPAR) inhibitors, cyclooxygenase (COX) inhibitors, angiotensin receptor inhibitors, vesicular monoamine transporter type (VMAT) inhibitors, retinoid acid receptor agonists, acetylcholine (ACh) effectors, copper chelators, cytomegalovirus (CMV) inhibitors, thrombopoietin receptor agonists, phosphodiesterase (PDE) inhibitors, 5α-reductase inhibitors, alkylating agents, and vitamin D agonists. Collectively, these 64 compounds comprise a heterogeneous mix of thoroughly studied and seldom explored potential therapeutic targets for neurodegenerative diseases. One of the main advantages of high-throughput behavioral screening approaches for drug repurposing is the lack of assumption regarding mechanism. In this study we present a diverse collection of compounds with seemingly diverging mechanisms of action, yet all evoking similar behavioral profiles in zebrafish larvae. Because AD is a multi-faceted neurodegenerative disease, it can greatly benefit from multi-targeted approaches to potentially ameliorate its pathology and impede further progression.

## Methods

### Animal Handling and Husbandry

All of the research in this study has been conducted in accordance with federal regulations and guidelines for the ethical and humane use of animals and has been reviewed and approved by Brown University’s Institutional Animal Care and Use Committee (IACUC). All behavioral experiments were performed on 5 days post-fertilization (dpf) zebrafish larvae (Danio rerio). Wild-type adult zebrafish used for breeding were housed at Brown University’s Animal Care facilities in 15- and 30-gallon tanks containing a mixed male and female population and kept on a 14 hr light and 10 hr dark cycle. During the light cycle, adult zebrafish were fed with Gemma Micro 300 and frozen brine shrimp. Adult zebrafish were bred in a group setting of approximately 40 zebrafish per tank, and embryos were collected and grown to 5 dpf as previously described(Pelkowski et al., 2011; Thorn et al., 2019; Tucker Edmister, Del Rosario Hernández, et al., 2022; Tucker Edmister, Ibrahim, et al., 2022). Embryos and larvae (0-5 dpf) were housed in 2L tanks with egg water containing 60 mg/L sea salt (Instant Ocean) and 0.25 mg/L methylene blue in deionized water. Zebrafish larvae used in this study were not fed, as they can obtain proper nutrition from their yolk sacs (Clift et al., 2014). Additionally, because sexual dimorphism is not apparent at this stage, larvae were not differentiated by sex (Liew & Orbán, 2014).

### Pharmacological Treatment

Zebrafish larvae were treated with 876 FDA-approved compounds from the Cayman Chemical FDA-Approved Drugs Screening Library (Cayman Chemical, Ann Arbor, Michigan, Item No. 23538). Each compound was originally provided in 10mM stocks dissolved in dimethyl sulfoxide (DMSO) and we diluted these compounds in egg water to a 10μM final concentration. Zebrafish larvae at 5 dpf were exposed to 100μL of the treatment or control solutions for a total of 6 hours. Egg water and 1μl/ml DMSO were used as control treatments.

### Behavioral Imaging

Zebrafish larvae were imaged using our previously described imaging setup and protocols (Pelkowski et al., 2011; Thorn et al., 2019; Tucker Edmister, Del Rosario Hernández, et al., 2022; Tucker Edmister, Ibrahim, et al., 2022). Briefly, larvae were imaged after 3 hours of exposure while placed in 96-well opaque plates (PerkinElmer, 6006290). The imaging system contains a glass stage capable of holding 4 plates at once. The plates were placed in a temperature-controlled imaging cabinet kept at 28.5℃. A high-resolution camera (18-megapixel Canon EOS Rebel T6 with an EF-S 55–250 mm f/4.0–5.6 IS zoom lens) captures a picture of the plates every 6 seconds. An M5 LED pico projector (Aaxa Technologies) with a 900 lumens LED light source was used to display a 3-hour Microsoft PowerPoint presentation featuring visual stimuli in the form of moving lines, as well as audio stimuli (100 ms, 400Hz) repeating at 1- and 20-second intervals(Gore et al., 2023).

### Image Analysis

Larvae pose estimation and subsequent behavioral quantification were performed using our automated image processing framework, Z-LaP Tracker, which contains a model trained with open-source software DeepLabCut(10, 66). Briefly, we trained a deep neural network to recognize three main features of zebrafish larvae: the right eye, left eye, and yolk sac. This model allows us to identify larvae even in changing background conditions. The quantified larval behaviors generated by Z-LaP Tracker were further evaluated and summarized using Excel templates (10). We evaluated a total of 25 behaviors encapsulating activity, reactivity, swimming patterns, and optomotor response (10). The differences of these values compared to DMSO-vehicle controls formed behavioral paradigms for each compound.

### Cluster Analyses

We evaluated a total of 876 compounds from the Cayman Chemical FDA-Approved Drugs Screening Library. We exposed 5 dpf zebrafish larvae to each compound (n=48 larvae) and averaged the results of each compound. Additionally, we normalized the data by calculating differences in behavior in comparison to the DMSO-vehicle control, creating behavioral profiles suitable for cluster analysis.

We performed K-means, a distance-based algorithm that seeks to partition each data point into one of k number of clusters by iteratively minimizing the distance between a point and its corresponding cluster mean. We used principal component analysis (PCA) as a dimensionality reduction method, and we used the elbow method to determine an optimal number of clusters k (k=4). PCA transforms the data into a two-dimensional form where K-means can be applied, and the clustering results plotted in a geometrical space.

We also performed hierarchical clustering, an unsupervised clustering method that allows us to visualize hierarchical relationships between individual compounds and their assigned clusters. Specifically, we used agglomerative hierarchical clustering with Euclidean distance as a distance metric, and complete linkage to measure dissimilarities between clusters.

All clustering methods were performed using R (R Studio 2022.12.0).

### Library Annotation and IPA Analysis

Biological targets and pathways were assigned to each of the compounds in the Cayman Chemicals FDA-approved drug library, based on hits from the Disease-Gene Interaction Database (DGIdb)(Freshour et al., 2021), Therapeutic Targets Database (TTD)(Chen et al., 2002), Guide to Pharmacology (GtoPdb)(Harding et al., 2022), Kyoto Encyclopedia of Genes (KEGG)(Kanehisa & Goto, 2000), Protein ANalysis THrough Evolutionary Relationships (PANTHER)(Mi & Thomas, 2009), WikiPathways(Slenter et al., 2018), and Reactome(Jassal et al., 2020) databases. Each database was queried for primary and secondary target genes associated with the compounds, as well as their respective molecular pathways and associated mechanisms of action. Cross-database datasets were generated with available matching information (i.e., UniProt ID, Gene Symbol, Ligand ID) to annotate the library. QIAGEN Ingenuity Pathway Analysis (IPA) was used to further analyze compounds of interest. Specifically, we used the “Build a Pathway” tool to input previously identified targets from our database query associated with compounds of interest. We then added “Alzheimer’s disease” and its related targets as a node in our pathway. Connections between elements were automatically generated using IPA’s “Connect” tool. We generated an overlay using the “Molecule Activity Predictor (MAP)” to indicate activation or inhibition of pathway components and connections.

### Statistical analyses

Statistical tests and graphs were generated using Microsoft Excel 2016, R, and BioRender. Due to the nature of our data, we used non-parametric Welch’s unequal variance t-test along with a Bonferroni correction for multiple comparisons. In the current screen, we compared 876 drugs to the DMSO-vehicle controls and differences were considered significant when p < 5.7 x 10-5 p < 1.1 x 10-5 (0.01/876), or p < 1.1 × 10-6 (0.001/876). Pearson correlation coefficients were calculated using R.

## Acknowledgments

We are grateful for valuable intellectual input from members of the Creton lab. This work was supported by the National Institutes of Health, Grant R01GM136906 (R.C.) and Grant R01GM136906-03S1 (R.C., J.A.K.).

## Author contributions

Conceptualization: RC, JAK Methodology: RC, SVG Investigation: TDRH, SVG Visualization: TDRH Supervision: RC, SVG, JAK Writing—original draft: TDRH Writing—review & editing: RC, SVG, TDRH

## Data Sharing and Availability

The DeepLabCut model used for pose estimation can be found on our GitHub repository (https://github.com/brown-ccv/Automated-Analysis-of-Zebrafish), along with installation and usage instructions. All data are available in the main text or the supplementary materials.

## Supplementary Materials

**Fig. S1. (A)** Subclusters of CsA-like compounds identified with hierarchical clustering. **(B)** Box-whisker plots of Pearson correlation values of compounds with the same primary target; mechanisms of action are marked (+) for agonists and (-) for antagonists. Pearson correlation values for the subclusters of CsA-like compounds are also plotted. The mean correlation of biologically unrelated compounds is plotted as a red line (0.186).

**Fig. S2.** Hierarchical cluster analysis of the behavioral profiles for all screened compounds.

**Fig. S3.** Behavioral profiles of 19 compounds previously identified as CsA-like during the screening of the Tocriscreen FDA-Approved Drugs Library. **(A)** 12 compounds displaying dissimilar behavioral effects from CsA. Behavioral profiles induced by biologically similar compounds, namely **(B)** β2-adrenergic receptor agonists, **(C)** α1-adrenergic receptor blockers, **(D)** androgen blockers, **(E)** PDE5 inhibitors, and **(F)** VEGFR inhibitors.

**Table S1.** Difference values of drug-treated larvae vs DMSO-treated larvae for all 876 compounds.

**Table S2.** Statistical analyses performed on the behavioral profiles of all 876 compounds.

