## Supplementary figures and images for "Finding Drug Repurposing Candidates for Neurodegenerative Diseases using Zebrafish Behavioral Profiles"

### Supplementary Figure 1

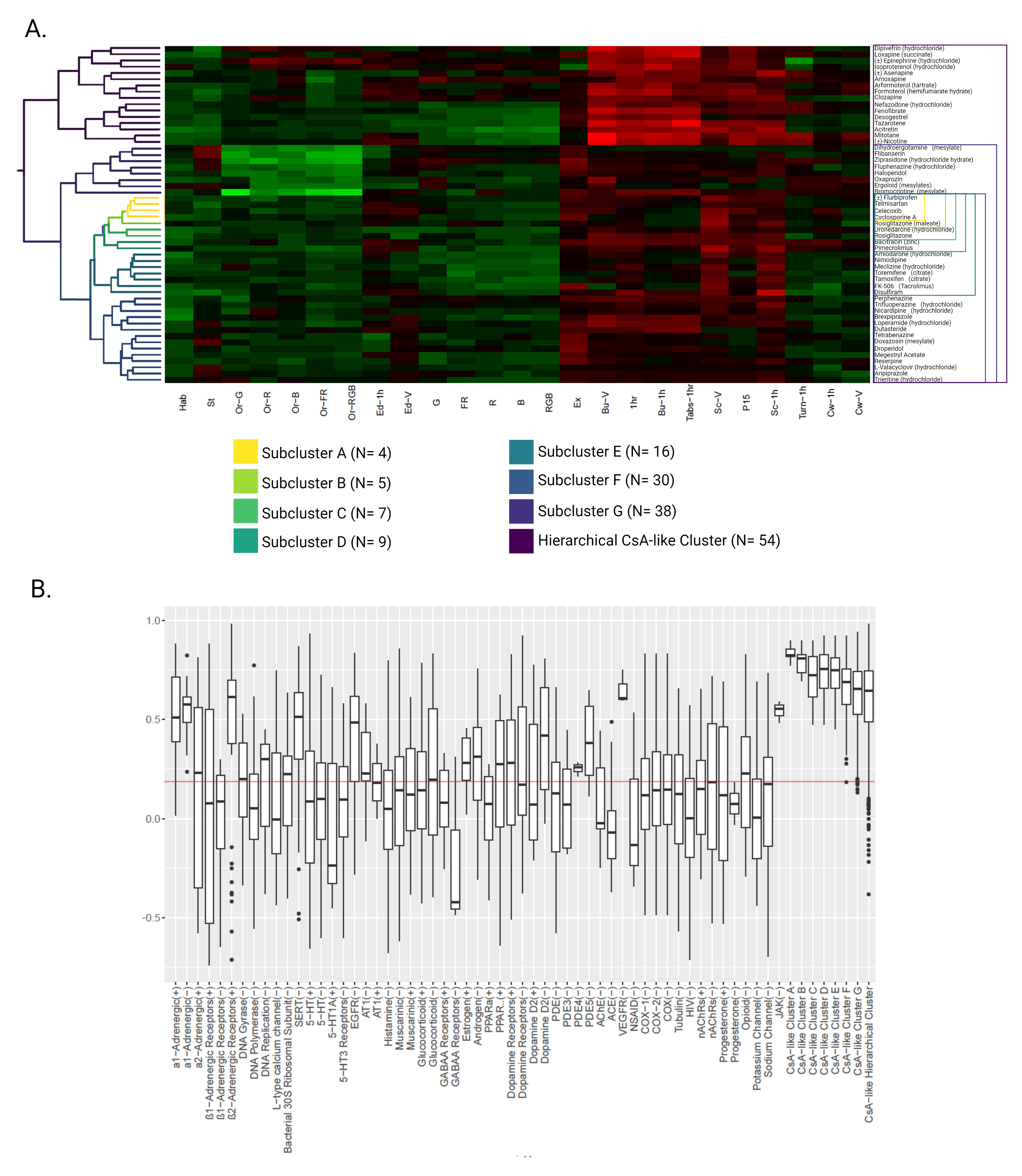

### Supplementary Figure 2

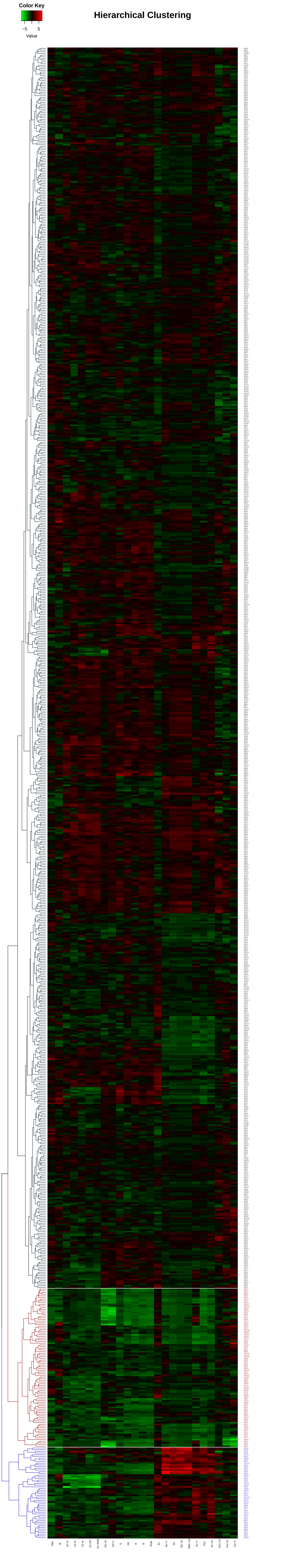

### Supplementary Figure 3

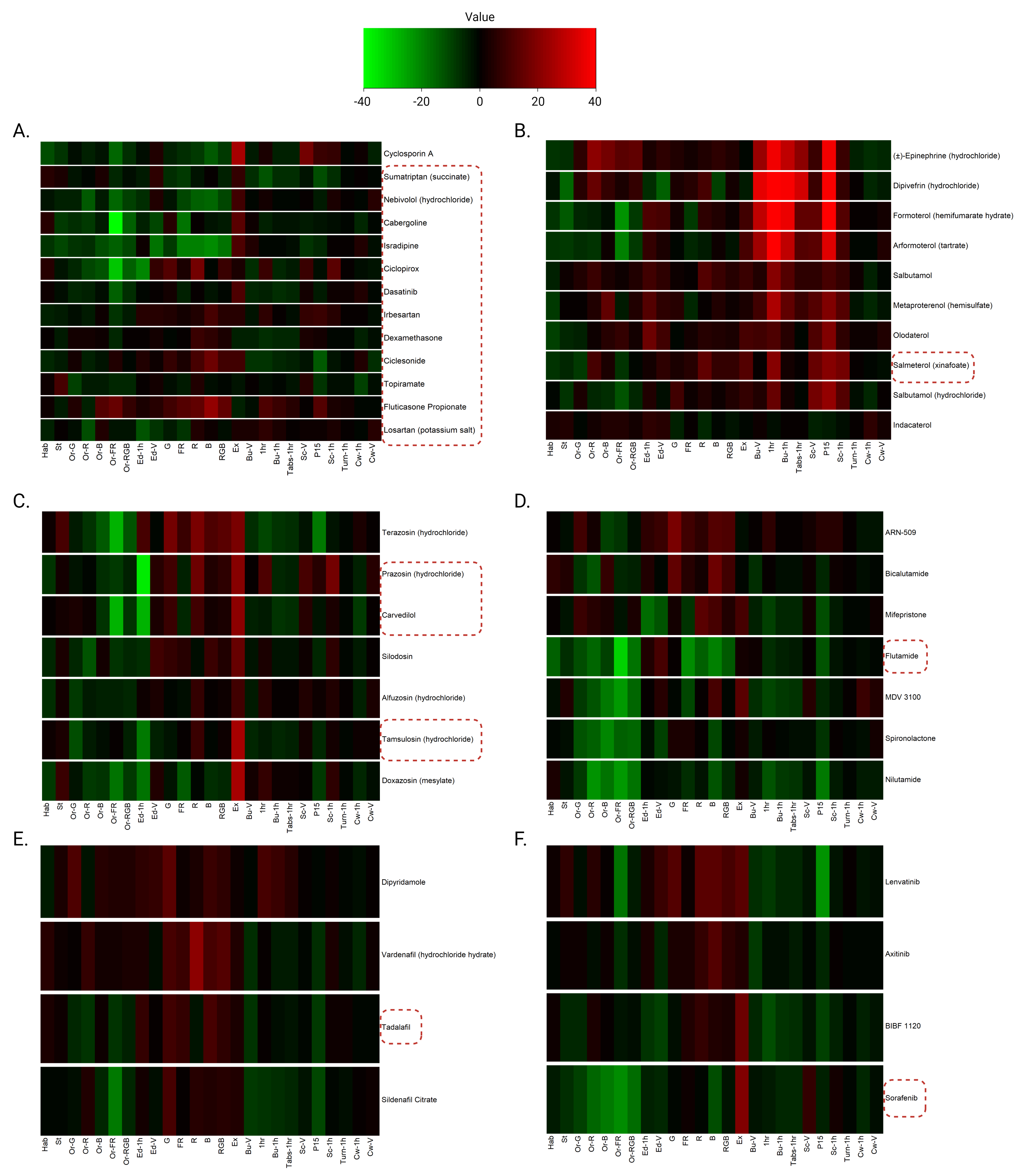
